# Decoupling Dopamine Synthesis from Impulsive Action, Risk-related Decision-Making, and Propensity to Cocaine Intake: A Longitudinal [^18^F]-FDOPA PET Study in Roman High- and Low-avoidance Rats

**DOI:** 10.1101/2023.11.29.569200

**Authors:** Ginna Urueña-Méndez, Chloé Arrondeau, Lidia Bellés, Nathalie Ginovart

**Affiliations:** Faculty of Medicine, Department of Psychiatry, University of Geneva, Geneva, Switzerland; Faculty of Medicine, Department of Basic Neuroscience, University of Geneva, Geneva, Switzerland

**Keywords:** [^18^F]-FDOPA, DA synthesis, Impulsive Action, Risk-Related Decision-Making, Cocaine SA

## Abstract

Impulsive action and risk-related decision-making (RDM) are two facets of impulsivity linked to a hyperdopaminergic release in the striatum and an increased propensity to cocaine intake. We previously showed that with repeated cocaine exposure, this initial hyperdopaminergic release is blunted in impulsive animals, potentially signaling drug-induced tolerance. Whether such dopaminergic dynamics involve changes in dopamine (DA) synthesis as a function of impulsivity is currently unknown. Here, we investigated the predictive value of DA synthesis for impulsive action, RDM, and the propensity to take cocaine in a rat model of vulnerability to cocaine abuse. Additionally, we assessed the effects of cocaine intake on these variables. Rats were tested sequentially in the rat Gambling Task (rGT) and were scanned with positron emission tomography and [^18^F]-FDOPA to respectively assess both impulsivity facets and striatal DA synthesis before and after cocaine self-administration (SA). Our results revealed that baseline striatal levels of DA synthesis did not predict impulsive action, RDM, or a greater propensity to cocaine self-administration (SA) in impulsive animals. Besides, we showed that impulsive action, but not RDM, predicted higher rates of cocaine-taking. However, chronic cocaine exposure had no impact on DA synthesis nor affected impulsive action and RDM. These findings indicate that the hyperresponsive DA system associated with impulsivity and a propensity for cocaine consumption, along with the reduction in this hyperresponsive DA state in impulsive animals with a history of cocaine use, is not mediated by dynamic changes in DA synthesis.

**Significance statement:** Impulsive behaviors are associated with a heightened presynaptic dopamine (DA) function and vulnerability to the rewarding effects of cocaine. However, with repeated drug exposure, the initially high DA release decreases, probably reflecting the development of drug tolerance. Whether such DA dynamics involve changes in DA synthesis is currently unknown. Using in vivo neuroimaging in rats before and after chronic cocaine use, our study reveals that DA synthesis does not predict impulsivity or vulnerability to cocaine, nor is it affected by chronic drug exposure. Our results suggest that the heightened presynaptic function underlying impulsivity and the cocaine-induced tolerance to drugs depend on alternative mechanisms to DA synthesis, such as those controlling DA reactivity to stimulation and DA reuptake.

## Introduction

Impulsive action and risk-related decision-making (RDM) are two facets of impulsivity differentially linked to individual vulnerability to cocaine abuse (Dalley and Robbins, 2017). While impulsive action predicts greater cocaine intake (Arrondeau et al., 2023; Dalley et al., 2007), even under punishment (Belin et al., 2008), RDM predicts the incubation of craving (Ferland and Winstanley, 2017). Despite their differential contribution to the risk of cocaine abuse, both impulsivity facets share dopaminergic alterations akin to those observed in drug abusers (Dalley and Robbins, 2017). For instance, humans with high trait impulsivity exhibit reductions in striatal dopamine (DA) D_2/3_ receptors (D_2/3_R) (Buckholtz et al., 2010) as those observed in cocaine abusers (Volkow et al., 1993). Such reduction in striatal D_2/3_R also occurs in rodents with high impulsive action (Bellés et al., 2021; Dalley et al., 2007) and RDM (Simon et al., 2011) and predicts a higher cocaine intake (Bellés et al., 2021; Dalley et al., 2007). Additionally, trait impulsivity and RDM are linked to heightened DA release in response to psychostimulants in humans (Buckholtz et al., 2010; Oswald et al., 2015), and such elevated DA release has also been confirmed in impulsive rats (Bellés et al., 2021). Importantly, heightened DA release predicts striatal D_2/3_R deficits, suggesting that those deficits might represent a compensation for heightened DA levels in impulsive animals (Bellés et al., 2021). Moreover, hyperdopaminergic release in impulsive animals predicts a higher propensity to cocaine self-administration (SA) (Urueña-Mendez et al., 2023). Together, these studies strengthen the view that increased presynaptic -rather than decreased postsynaptic-DA function might be the main dysregulation underlying impulsive behaviors, conferring greater vulnerability to cocaine abuse.

Although the molecular underpinnings of such heightened DA function in impulsive subjects are unknown, one potential underlying mechanism could be increased DA synthesis. Indeed, tyrosine hydroxylase (TH) expression is higher in the midbrain of innately high-impulsive rats relative to low-impulsive rats (Tournier et al., 2013). However, no study has yet evaluated whether impulsive action and RDM are associated with alterations in DA synthesis and how such alterations might predict the propensity to take cocaine.

Besides the predictive value of impulsivity facets and DA function in cocaine abuse, there is also evidence that chronic cocaine exposure can affect impulsive behaviors and DA signaling (Dalley and Robbins, 2017). However, cocaine effects may vary depending on the impulsivity facet assessed. While several studies (Abbott et al., 2022; Dalley et al., 2005; Ferland and Winstanley, 2017; Urueña-Mendez et al., 2023) have reported no alterations in impulsive action shortly after cocaine withdrawal (but see: Dalley et al., 2007; Winstanley et al., 2009), research on RDM is scarce and has yielded mixed results. For instance, one study (Ferland and Winstanley, 2017) found impaired RDM exclusively in risky rats (i.e., those preferring high reward magnitude at higher punishment risk), while another study found impairments in non-risky rats (Cocker et al., 2019). Such inconsistencies highlight the need for further longitudinal studies to clarify the consequences of chronic cocaine use on different impulsivity facets. Chronic cocaine use also leads to neuroadaptive alterations in DA signaling. Indeed, chronic cocaine exposure attenuates DA release in response to psychostimulants in impulsive rats (Urueña-Mendez et al., 2023), which is consistent with the blunted DA release observed in cocaine abusers (Martinez et al., 2007; Volkow et al., 2014). Although the mechanisms underlying such dopaminergic tolerance to psychostimulants are unclear, one study on cocaine abusers suggests that it could involve reductions in DA synthesis (Wu et al., 1999). However, the study’s results have been debated (Cumming et al., 1999), and no additional research on cocaine abusers has been performed to confirm this finding. Similarly, no preclinical studies have ever assessed whether chronic cocaine use affects DA synthesis in relation to impulsivity. Altogether, current research links impulsive action and RDM to greater DA release and higher risk of cocaine abuse, and also indicates that chronic cocaine use can induce DA tolerance to psychostimulants. However, whether such DA dynamics involve changes in DA synthesis is currently unknown.

Here, we used in vivo positron emission tomography (PET) and the L-DOPA radiolabelled analog ^18^F-fluoro-dihydroxyphenylalanine ([^18^F]-FDOPA) to investigate the predictive value of striatal DA synthesis on impulsive action, RDM, and propensity to cocaine SA in the roman high-(RHA) and low-avoidance (RLA) rat model of vulnerability to drug abuse (Giorgi et al., 2019). Additionally, we evaluated the effects of chronic cocaine intake on impulsive behaviors and DA synthesis. Our results shed light on the DA mechanisms underlying impulsivity and vulnerability to the rewarding effects of cocaine.

## Materials and methods

### Subjects

Subjects were outbred male rats RLA (n=22) and RHA (n=23) from our permanent colony at the University of Geneva. Rats were paired or trio housed in a temperature-controlled room (22±2°C) under a 12h light-dark cycle (lights on at 07:00 a.m.). Food was restricted daily (5g per 100g of body weight) throughout the experiment, and water was provided ad libitum. All experiments were approved by the animal ethics committee of the Canton of Geneva and performed according to the Swiss federal law on animal care.

### General procedure

Figure 1 shows the timeline of the study. Two cohorts of rats (n=22-23 per line) were first trained and tested in the rat Gambling Task (rGT) to concurrently measure their baseline levels of impulsive action and RDM. Next, a subgroup of rats in each cohort (16 rats per line) was scanned with PET and [^18^F]-FDOPA to assess their baseline capacity to synthesize DA. Rats were then trained to self-administer either cocaine (n=14-15 rats per line) or saline vehicle (n = 8 rats per line). Three days after the last SA session, the subgroup of rats scanned at baseline received a second [^18^F]-FDOPA PET scan. Finally, starting at day 6 post-SA, all rats were re-tested in the rGT for 8 days. This procedure allows us to investigate the effects of repeated cocaine exposure on DA synthesis capacity, impulsive action, and RDM.

**Figure 1.**
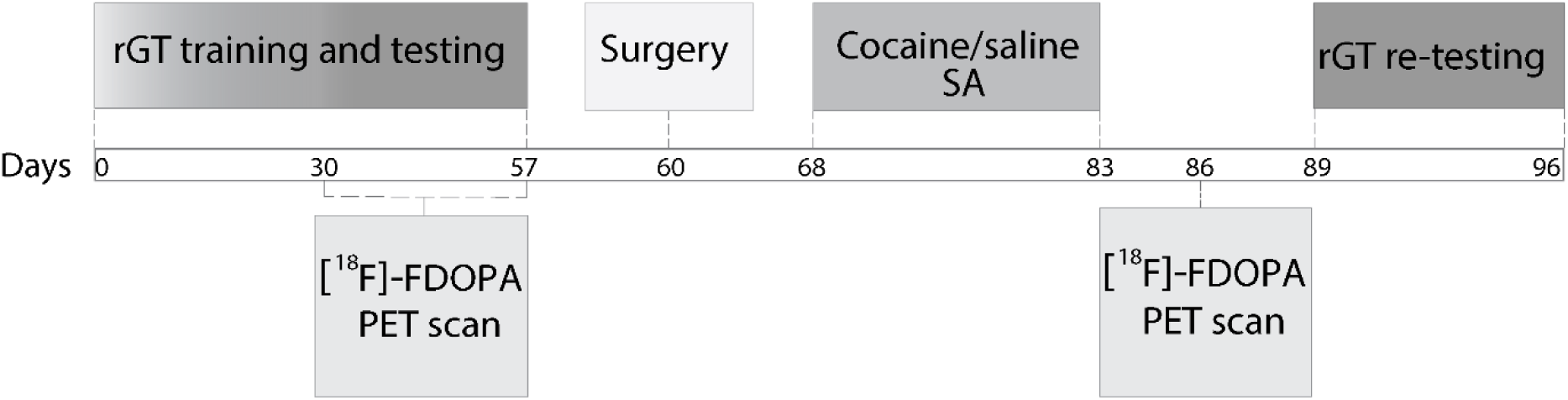
Timeline of the study with specific time points (in days) annotated in the main line.

### Rat Gambling Task (rGT)

The rGT task was performed in eleven standard operant conditioning chambers (Med Associates Inc., St. Albans, VT, USA). Each chamber was fitted with a house light and a food magazine. On the opposite wall, five response holes (the central hole being inactivated during the task) were fitted with a cue light and infrared beams to detect nose-poke responses.

Testing occurred daily (5d/week) in 30min sessions performed between 8-12h following a protocol previously described (Zeeb et al., 2009). In brief, animals learned to nose-poke in one illuminated (i.e., active) hole to obtain a food pellet (purified rodent tablets of 45 mg, Test Diet, Sandown Scientific, UK). Subsequently, rGT options were introduced over seven forced-choice sessions. The options were termed P1, P2, P3, and P4. They varied in the number of pellets delivered (1, 2, 3, or 4, respectively), the probability of pellet delivery (0.9, 0.8, 0.5, or 0.4, respectively), the time-out punishment duration (5s, 10s, 30s, or 40s, respectively) and the time-out punishment probability (0.1, 0.2, 0.5, or 0.6, respectively) for the non-rewarded trials. The options were associated with specific holes counterbalanced between animals.

Rats initiated each trial by nose-poking into the food magazine. This response triggered a 5s intertrial interval (ITI). Once the ITI had elapsed, one option was pseudorandomly presented (i.e., the corresponding hole lit), and the rat had 10s to nose-poke. This response was followed by reward or punishment according to the hole-associated contingency. During punished trials, the hole light flashed at 0.5Hz. After completing the forced-choice training, rats performed the rGT task (i.e., with all the options presented simultaneously) during 25 sessions to establish a stable response at baseline. A choice score [(%P1+%P2)–(%P3+%P4)] was used as an index of RDM (Fugariu et al., 2020; Hultman et al., 2022; Zeeb and Winstanley, 2013), with a lower choice score indicating less optimal decision-making. Impulsive action was defined as the percentage of premature responses performed during the ITI [i.e., number of premature responses/ total number of trials initiated x 100] (Ferland and Winstanley, 2017). Both outcome measures were calculated at baseline and after cocaine SA as the average across three consecutive sessions of stable performance, with no main effect of session and no session-by-choice score interaction as assessed using a repeated-measures analysis of variance (ANOVA).

### [^18^F]-FDOPA PET imaging

Animals were fasted for 12h before the PET scan. To prevent early breakdown and decarboxylation of [^18^F]-FDOPA (Firnau et al., 1986; Sawle et al., 1994), rats received 40 mg/kg (ip) of entacapone (Combi-Blocks, Switzerland) 1h before the scan and 10 mg/kg (ip) benserazide hydrochloride (Sigma-Aldrich, Switzerland) 30min before the scan, respectively. Under isoflurane anaesthesia, two animals per scan were positioned in the micro-PET scanner Triumph II (TriFoil Imaging, Northridge, CA, USA) using a custom-made stereotactic-like frame. After a 5-minute computer tomography (CT) scan, rats received an i.v. bolus injection of 58±9 MBq of [^18^F]-FDOPA (Advanced Accelerator Applications, Saint-Genis-Pouilly, France) and were scanned for 60min.

Dynamic PET images were acquired in list mode and reconstructed into 19 timeframes (2 x 30s, 2 x 60s, 4 x 120s, 3x 180s, 8 x 300s) using the ordered subsets expectation maximization (OSEM) algorithm with 30 iterations. Image analysis was conducted with PMOD software (version 4.0, PMOD Technologies Ltd., Zurich, Switzerland). PET summation images were generated from dynamic PET data and checked for alignment with the corresponding CT images. Because minor motion occurred between the PET and CT images, all PET images were coregistered to the corresponding CT images using rigid registration based on the normalized mutual information. CT images were then coregistered to a magnetic resonance image (MRI) brain template of the rat brain (Schiffer et al., 2006) using mutual information-based rigid body registration. The resulting transformation was then applied to the corresponding PET dynamic images, mapping all rat brains into the same reference space.

A region of interest (ROI) template was defined on the MRI atlas of the rat brain (Schweinhardt et al., 2003), as previously described (Ginovart et al., 2012). This template included three brain regions: the dorsolateral striatum (DST), the ventral striatum (VST), and the cerebellar cortex. ROIs (2mm diameter circles) for the DST and VST were placed on the central planes of each structure to minimize the partial volume effect. A single elliptical ROI (2mm x 1mm) was placed on the cerebellar cortex, primarily over the central lobule, to reduce spillover from bone uptake of [^18^F]-fluoride. The ROI template was applied to the dynamic images to produce time-activity curves for the target-rich (DST) and reference region (cerebellar cortex), which were then used to generate parametric maps. Voxel-wise parametric maps of the [^18^F]-FDOPA influx rate constant (*k_i_^cer^*, min^-1^) were calculated using the Patlak multiple-time graphical analysis (Patlak and Blasberg, 1985) using the cerebellum as a reference to estimate the free and non-specifically bound radiotracer. The ROI template was then applied to the individual parametric maps to extract regional *k_i_^cer^* estimates as an index of DA synthesis. Both hemisphere data were averaged to obtain a single *k_i_^cer^* value for the DST and VST.

### Cocaine self-administration (SA)

The SA procedure was performed as described in Dimiziani et al. (2019). In brief, rats were implanted with a polyurethane catheter (Instech Laboratories, Plymouth Meeting, PA, USA) into the right jugular vein under 2.0% isoflurane anesthesia. The catheter was fixed in the mid-scapular region and attached to a vascular access button (VAB) protected from cage mate chewing with an aluminum cap (Instech Laboratories, Plymouth Meeting, PA, USA). Rats received post-operatory care and were allowed to recover for one week before SA training. The catheter’s patency was maintained by infusing 0.1 ml of heparinized saline (30U.I./ml) daily. Training occurred in eleven standard operant conditioning chambers (Med Associates Inc., St. Albans, VT, USA) equipped with a house light and two response holes connected to an external infusion pump. The holes were fitted with a cue light and infrared beams to detect nose-poke responses. Rats were trained to self-administer 0.3 mg/kg/infusion of cocaine hydrochloride (Geneva, University Hospitals) or 0.9% saline vehicle under a fixed ratio 1 schedule (FR1). Sessions were performed daily (5 days/week) for three weeks and lasted 2h or until the animals reached the maximum 60 infusions allowed. Each session started with the illumination of the house light and a priming infusion of the assigned treatment (cocaine or saline). Nose-poke responses in a specific hole (active) extinguished the house light and turned on the hole-cue light while delivering one infusion of 0.06–0.1 ml of cocaine or saline over 2.5–4s. At the end of each infusion, the hole-cue light extinguished, signaling the beginning of a 20s time-out (TO). Nose-poke responses during the TO had no programmed consequences. Responses in the other hole (inactive) were recorded but delivered no infusions. The location of the active and inactive holes was randomly counterbalanced between animals.

Cocaine SA acquisition was defined as the obtention of ≥15 infusions during two consecutive days with accurate discrimination (70%) of the active vs inactive hole. Cocaine maintenance was defined as a stable drug intake (≤15% variation in the number of infusions) over three consecutive days. We evaluated the total number of infusions and rate of infusions as measures of drug intake and the total nose-poke responses in the active vs. inactive hole as a measure of hole discrimination/preference. For correlation analysis, the three last SA sessions were averaged to obtain a single value for each variable.

### Statistical analysis

All statistical analyses were conducted with SPSS Statistics 26.0 (IBM) software, except the *Grubbs*’ *Test* (at p<0.01) for outlier detection performed in GraphPad Prism 9.0 (GraphPad Software 9.0, San Diego, USA). Normality of data distribution was verified using the Shapiro-Wilk statistic at p<0.05, and non-normal data were LOG-10 transformed before the ANOVA analysis, as specified below. All ANOVA analyses were followed by Bonferroni post-hoc tests when appropriate.

For baseline comparisons, rats were grouped by line (RHA or RLA), irrespective of their subsequent treatment (saline or cocaine) assignment. Between-line differences in the percentage of premature responses and choice score were analyzed with an unpaired T-test or Man-Whitney-U statistic, depending on the data distribution. Baseline differences in [^18^F]-FDOPA k_i_ ^cer^ values were assessed using a two-way repeated measures ANOVA with line as between-subjects factor and brain region (DST or VST) as within-subjects factor.

The percentage of animals reaching the criteria for cocaine SA acquisition was evaluated using the Kaplan–Meier survival analysis. The discrimination of the active versus inactive hole during cocaine acquisition and maintenance was evaluated using a mixed factorial ANOVA, with session (days 1-15) and hole (active or inactive) as within-subjects factors and with line and treatment as between-subjects factors. In the measure of total nose-poke responses in the active hole, *Grubbs’ Test* identified one RLA outlier within the cocaine group (G=3.3 p<0.01). We applied the Winsorize method to correct for this outlier value in the ANOVA analysis.

The number of infusions and the rate of infusions displayed during acquisition and maintenance of cocaine SA were analyzed using a mixed factorial ANOVA, with session as within-subjects factor, and line and treatment as a between-subjects factors.

The effect of repeated cocaine exposure on premature responding, choice score and [^18^F]- FDOPA *k_i_^cer^* was evaluated using a mixed factorial ANOVA, with time (i.e., Pre- or Post-SA) as within-subjects factor, and line and treatment as between-subjects factors. Due to a technical failure in [^18^F]-FDOPA radiopharmaceutical synthesis, 1 RHA and 2 RLA rats did not receive their PET scan following SA and were therefore not included in the analysis.

Finally, correlations between [^18^F]-FDOPA *k_i_^cer^*, rGT, and SA variables were tested using Pearson’s or Spearman’s correlation according to the data distribution.

## Results

### Baseline Impulsive action and risk-related risky decision-making (RDM)

At baseline, RHA rats reached a significantly higher percentage of premature responses relative to RLAs (t=7.7, df=43, p<.001, d=2.3; Fig. 2A), indicating that RHAs displayed higher levels of impulsive action than RLAs. When evaluating RDM, RHAs displayed significantly lower choice scores than RLAs (U=166, p<.05, η^2^=.14; Fig. 2B), indicating less optimal decision-making than RLAs. No difference occurred in the total number of trials initiated by each rat line (t=-.7, df=43, p=.49 d=.20), thus showing that the between-line difference in choice score and premature responding did not result from differences in the trials performed by the two rat lines in the rGT.

**Figure 2.**
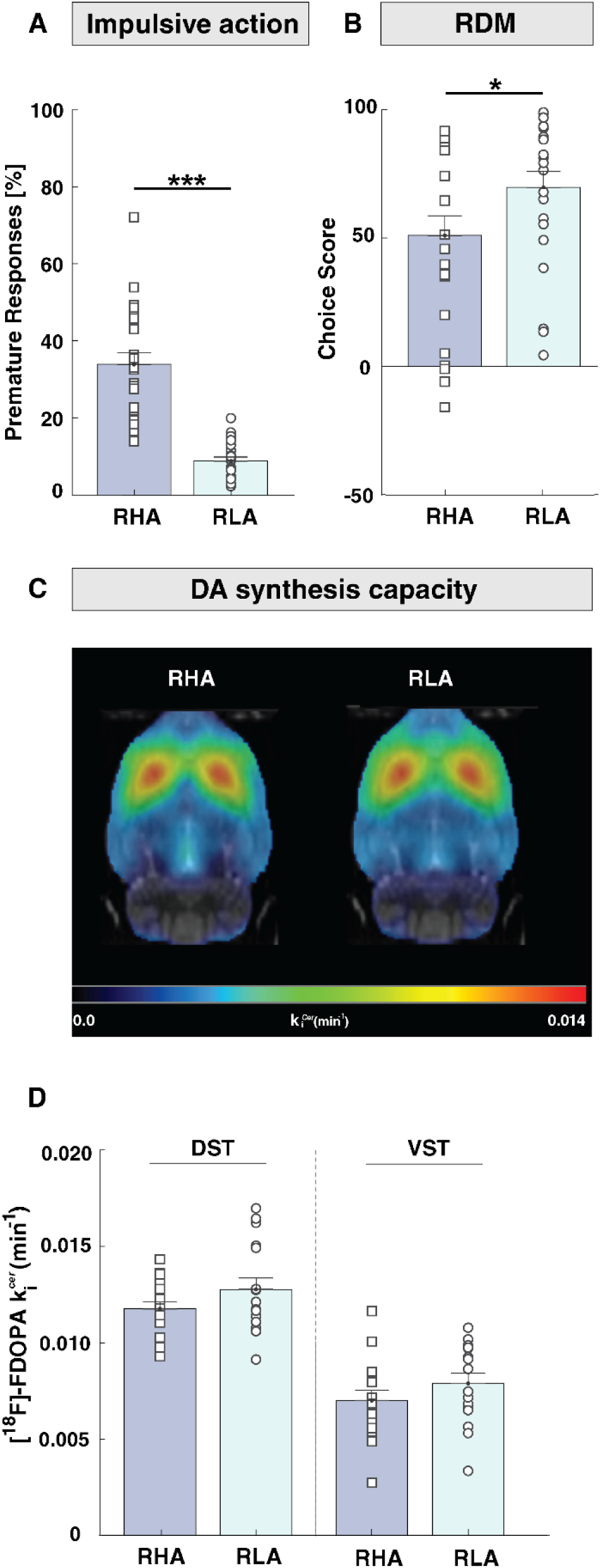
Baseline measures of impulsive action, RDM, and striatal DA synthesis capacity in RHA and RLA rats. **A)** The percentage of premature responses, as an index of impulsive action, was significantly higher in RHAs compared to RLAs. **B)** Choice score, as an index of RDM, was significantly lower in RHAs compared to RLAs. **C)** Mean parametric maps of [^18^F]- FDOPA k_i_ ^cer^ in RHA and RLA rats. k_i_ ^cer^ parametric maps are projected upon the MRI rat atlas (greyscale) and are shown in horizontal planes at the level of the DST. **D)** [^18^F]-FDOPA k_i_ ^cer^ did not differ between RHA and RLA rats in either DST or VST. Data are shown as mean ± SEM. Significantly different at *p<.05 and *** p<.001 in RHAs vs RLAs.

### Baseline striatal DA synthesis and impulsive behaviors

Figure 2C shows the mean parametric maps for [^18^F]-FDOPA *k_i_ ^cer^* obtained in RHA and RLA rats at baseline conditions. In both rat lines, [^18^F]-FDOPA *k_i_ ^cer^* values were higher in the DST than in the VST (Region: F_(1,30)_=150,9 p<.001, ηp^2^=.83; Fig. 2D). However, we found no significant differences in [^18^F]-FDOPA *k_i_^cer^* between the two rat lines (Line: F_(1,30)_ =2.38, p=.13, ηp^2^=.06) in either striatal subdivision (Line x Region: F_(1,30)_=.18, p=.89 ηp^2^=.001).

At baseline, there was no correlation between premature responding and [^18^F]-FDOPA *k_i_ ^cer^*in either the DST (r_(30)_=-.15, p=.40; Fig. 3A) or the VST (r_(30)_=-.05, p=.70; Fig. 3B), indicating that striatal [^18^F]-FDOPA *k_i_^cer^* does not predict impulsive action. Similarly, no correlation existed between the choice score and the baseline [^18^F]-FDOPA k*_i_^cer^* in any striatal subdivision *(DST:* r_(30)_=-.12, p=.50; *VST*: r_(30)_=-.06, p=.75; Fig. 3C**-D**), indicating that striatal [^18^F]-FDOPA *k_i_^cer^* is also not predictive of RDM.

**Figure 3.**
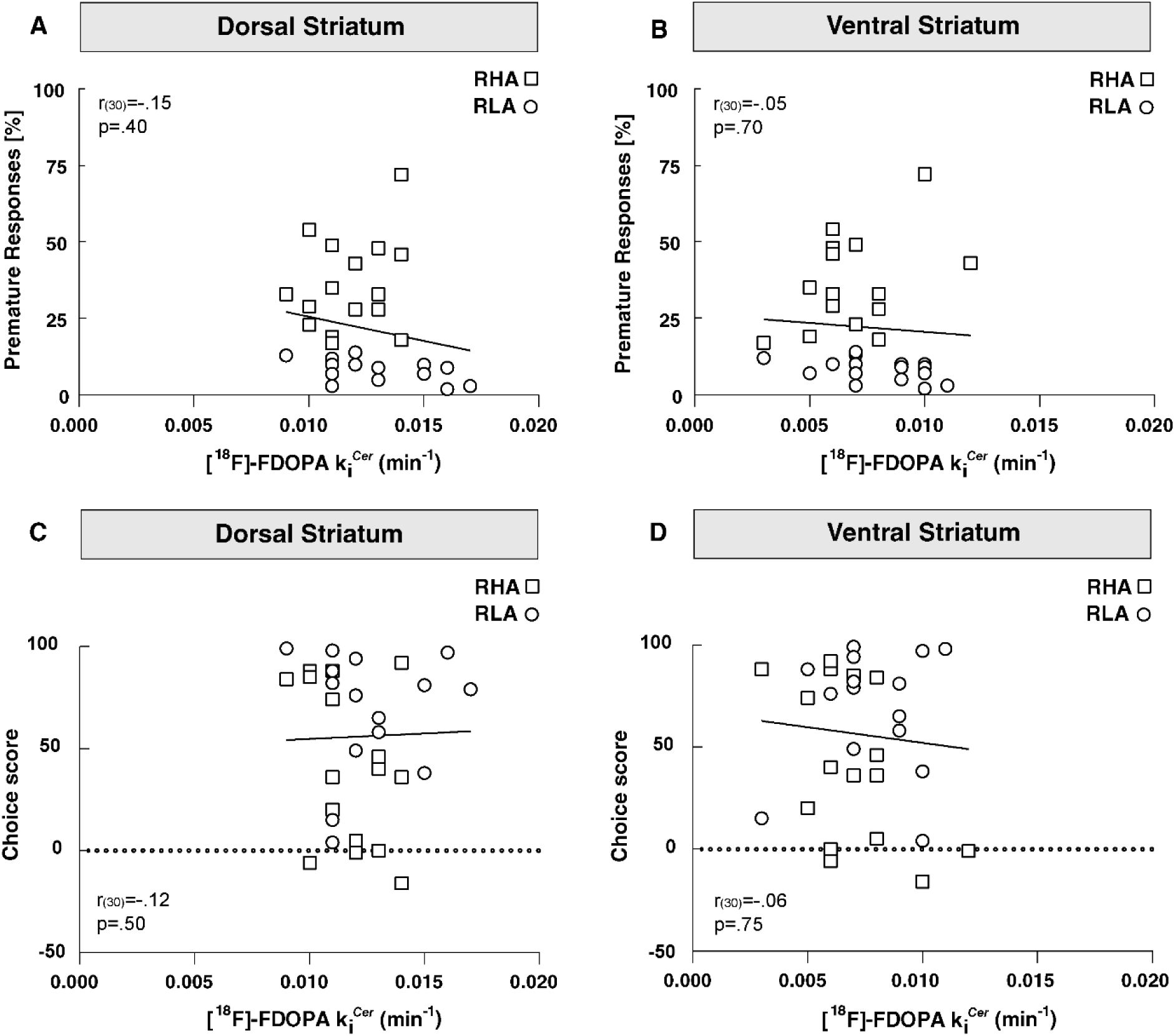
Relationships between striatal [^18^F]-FDOPA k_i_^cer^, impulsive action, and RDM in RHAs (□) and RLAs (○) at baseline. **A-B)** There was no correlation between [^18^F]-FDOPA k_i_^cer^ and premature responding in either the DST or VST. **C-D)** No correlation existed between [^18^F]-FDOPA k_i_^cer^ and choice score in either the DST or VST.

### Intravenous Cocaine Self-Administration

Figure 4 shows the acquisition and maintenance of cocaine and saline SA in RHA and RLA rats. SA acquisition (i.e., >15 infusions with 70% discrimination of the active hole over two consecutive days) occurred only in cocaine-exposed rats. In these rats, cocaine SA acquisition was faster in RHAs than in RLAs (χ^2^= 4.3, df=1, p>.05 Fig. 4A), with 80% of the RHAs reaching the acquisition criteria on day 2, while the same proportion of RLAs reached those criteria by day 5.

**Figure 4.**
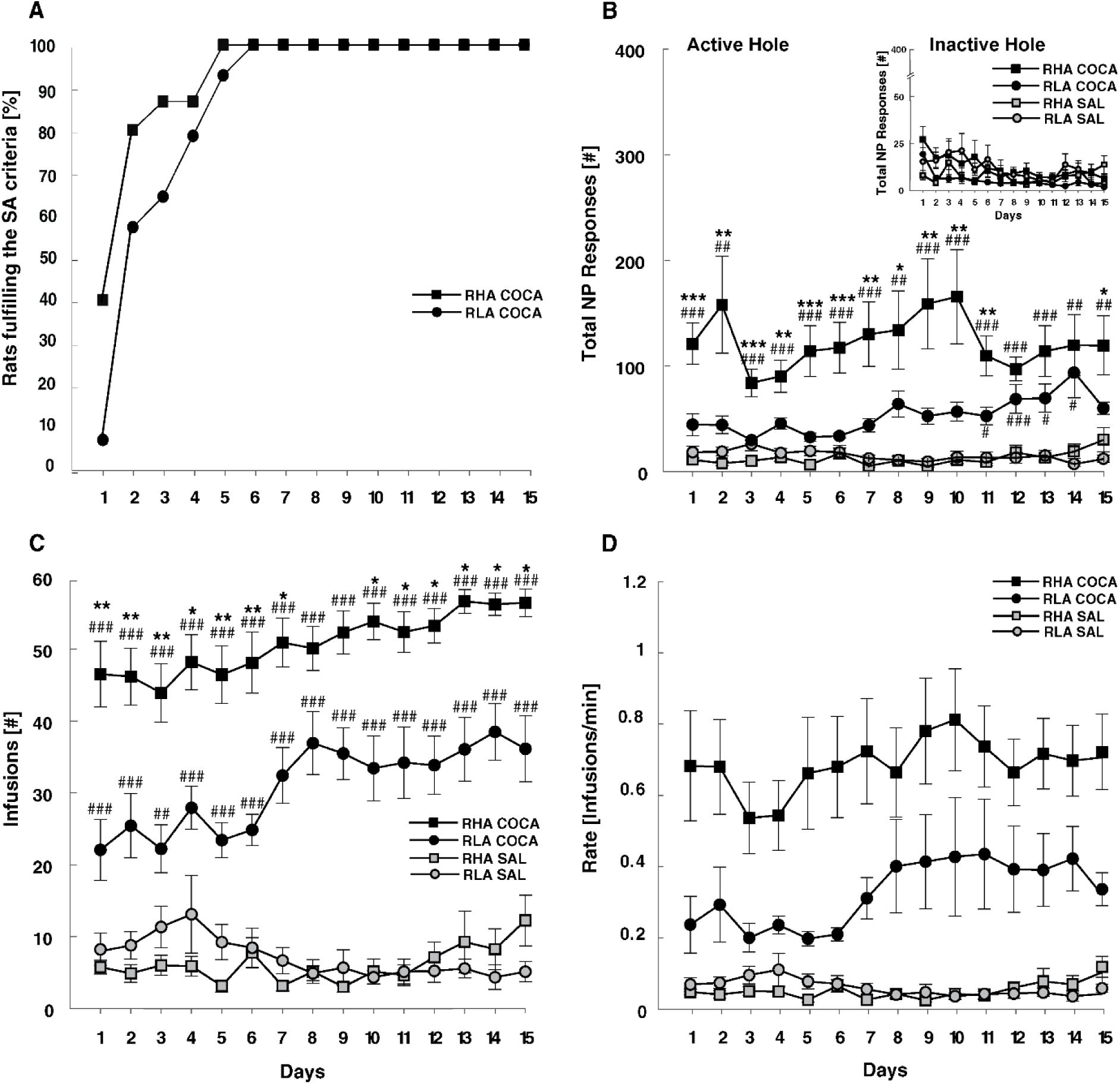
Comparison of cocaine and saline SA between RHA and RLA rats. **A)** Cumulative percentage of rats meeting SA acquisition criteria (i.e., >15 infusions with 70% discrimination of the active vs inactive hole over two consecutive days). Only cocaine-exposed animals fulfilled the SA acquisition criteria, with RHAs acquiring cocaine SA significantly faster than RLAs. **B)** Total nose-poke (NP) responses in the active hole were significantly higher in cocaine-than in saline-exposed rats and in RHAs compared to RLAs. The inserted image shows the responses in the inactive hole, which were similar between groups. **C)** The number of infusions was significantly higher in cocaine-exposed rats than saline-exposed rats, and in RHAs compared to RLAs. **D)** The rate of cocaine infusions was significantly higher in RHA than in RLA rats throughout the 15 days of SA. Data appears as mean ± SEM. Significantly different at * p<.05, ** p<.01 in cocaine-exposed RHAs compared to cocaine-exposed RLAs, and ## p<.01, ### p<.001 between cocaine-exposed RHAs or RLAs relative to their corresponding saline control.

A mixed factorial ANOVA on the number of nose-pokes in the active vs inactive hole (Fig. 4B) showed a main effect of Treatment (F_(1,42)_= 5.05, p<.001, ηp^2^=.34), Line (F_(1,42)_=5.05, p<.05, ηp^2^=.11), Hole (F_(1,588)_=37.6, p<.001, ηp^2^=.47), and Treatment x Line x Hole interaction (F_(1,588)_=5.14, p<.05, ηp^2^=.11) with no effect of Session (F_(1,588)_= 0.5, p=.93, ηp^2^=.01). Post-hoc comparisons further confirmed that only cocaine-exposed rats showed clear discrimination of the active vs inactive hole (RHAs p<.001; RLAs p<.001). Furthermore, cocaine-exposed RHAs performed statistically a higher number of nose-poke responses in the active hole than cocaine-exposed RLAs (p<.05). The number of infusions was also conditional to Treatment (F_(1,42)_=406.7 p<.001, ηp^2^=.91) and rat’s Line (F_(1,42)_=5.6 p<.05, ηp^2^=.12), with RHAs self-injecting more cocaine than RLAs along the training (Treatment x Line x Session: F_(14,588)_=2.8, p=.006 ηp^2^=.06; post hoc comparisons in RHA-Coca vs RLA-Coca, p<.05 except for days 8 and 9). Maintenance of cocaine SA (i.e., ≤15% variation in the number of infusions over three consecutive days) was also different between rat lines, with RHAs reaching the criteria of stable cocaine infusions on day 3, while RLAs reached the criteria on day 9 (Fig. 4C). Besides differences in cocaine-taking, RHAs, and RLAs also differed in how fast they self-injected cocaine (Fig. 4D). A mixed factorial ANOVA on the rate of cocaine infusion showed a main effect of Treatment (F_(1,42)_=31.3, p<.001, ηp^2^=.43), Line (F_(1,42)_=4.7, p<.05, ηp^2^=.10) and a Treatment x Line interaction (F_(1,42)_= 5.22, p<.05, ηp^2^=.11), with no effect of Session (F_(14,32)_=.80, p=.67, ηp^2^=.01). Post-hoc comparisons confirmed that cocaine-exposed RHAs were faster than cocaine-exposed RLAs to self-inject the drug (P<.05).

### Predictive value of impulsivity measures and DA synthesis on cocaine SA

There was a positive correlation between premature responding and the total number of infusions earned (r_(27)_=.6 p<.001, Fig. 5A). However, this correlation was potentially biased due to a ceiling effect resulting from the capped limit of 60 cocaine infusions allowed. Consequently, further analyses were conducted with the rate of cocaine infusions as a measure of cocaine-taking. We observed a positive correlation between premature responding and the rate of cocaine infusions, indicating that the higher the premature responses, the faster the animals self-injected cocaine (r_(27)_=.5 p<.001, Fig. 5B). No correlation was observed between premature responding and the total number of nose-poke responses in the active hole (r_(27)_=.3 p=.1). On the other hand, when evaluating the predictive value of the choice score, no correlation was found with the rate of cocaine infusions (r_(27)_=-.04 p=.81; Fig. 5C) or the total exploration of the active hole (r_(27)_=.03 p=.8). These results indicated that impulsive action but not RDM predicts the propensity to cocaine-taking.

**Figure 5.**
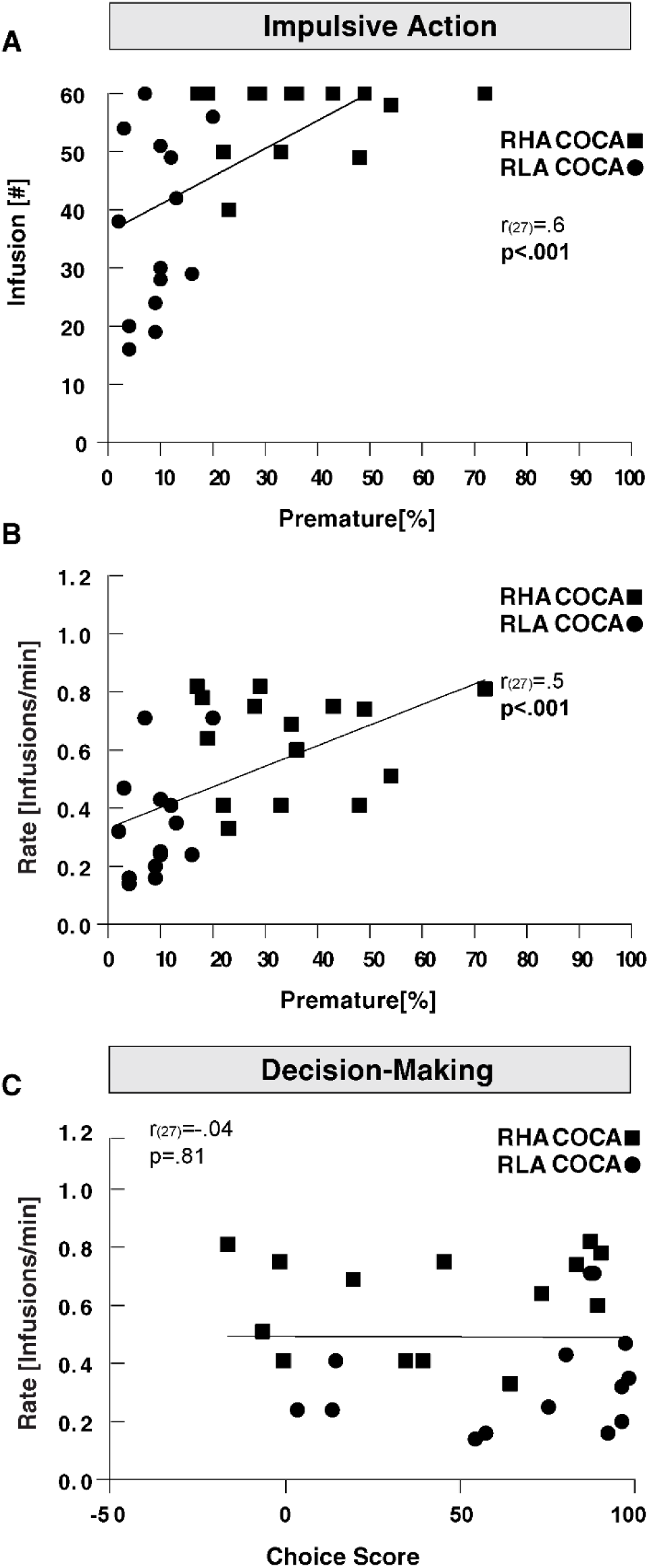
Relationship between impulsive action, RDM, and cocaine SA in RHA (▪) and RLA rats (●). Premature responding was significantly correlated with: A) the number of cocaine infusions earned, and B) the rate of cocaine infusions. C) No correlation was found between the choice score and the rate of cocaine infusions.

No correlation appeared between [^18^F]-FDOPA *k_i_ ^cer^* and the rate of cocaine infusions (DST: r_(17)_=-.29, p=.21; VST: r_(17)_=-.23 p=.32, Fig. 6**. A-B**), indicating that DA synthesis capacity does not predict cocaine-taking.

**Figure 6.**
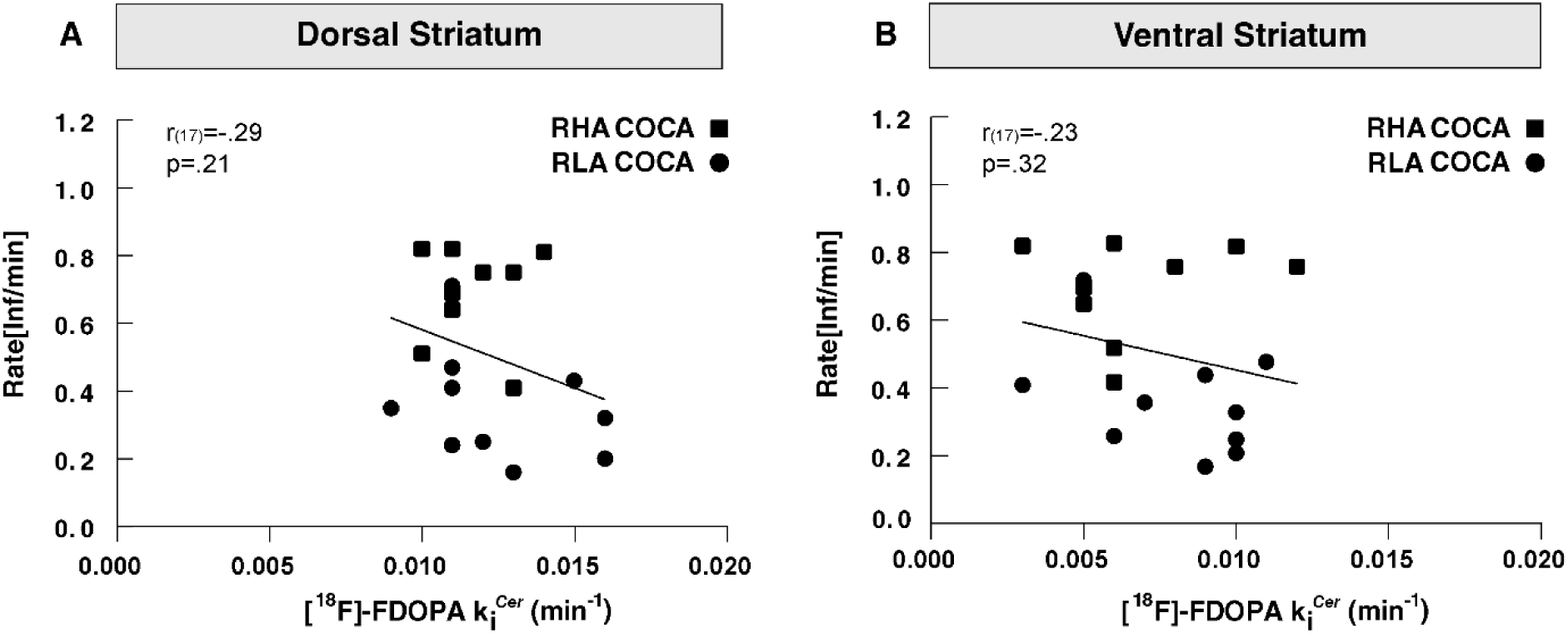
Relationship between [^18^F]-FDOPA k_i_^cer^ and cocaine SA in RHA (▪) and RLA rats (●). [^18^F]-FDOPA k_i_^cer^ in neither **A)** DST nor **B)** VST predicted the rate of cocaine infusions.

### Effect of repeated cocaine consumption on impulsive action and RDM

Figure 7 (A-B) shows the effect of prior cocaine or saline SA on rGT behaviors. Neither saline nor cocaine SA affected the percentage of premature responses in RHA or RLA rats (Treatment: F_(1,41)_=.22 p=.64, ηp^2^=.005; Treatment x Line: F_(1,41)_=.82 p=.37, ηp^2^=.02). Similarly, there was no effect of Time (F_(1,41)_ =.48 p=.49, ηp^2^=.012) or Time x Line x Treatment interaction (F_(1,41)_ =3.6 p=.064, ηp^2^=.08). These results indicate that baseline impulsive action is not significantly altered after cocaine or saline SA. Similarly, a mixed factorial ANOVA showed no effects of Treatment (F_(1,41)_ =.48 p=.49, ηp^2^=.12) or a Treatment x Line interaction (F_(1,41)_=1.29 p=2.6, ηp^2^=.03) on choice score. Further, there was neither an effect of Time (F_(1,41)_= 3.89, p=0.06, ηp^2^=0.08) nor a Time x Line x Treatment interaction (F_(1,41)_= 001 p=.97, ηp^2^=0) on choice score, indicating that cocaine or saline SA in both rat lines did not significantly alter RDM when compared to baseline. Finally, no changes in the number of trials occurred as a consequence of the Treatment (F_(1,41)_=.3, p=.87, ηp^2^=.001), Treatment x Line (F_(1,41)_=.43, p=.51, ηp^2^=.01), Time (F_(1,41)_= 2.06, p=.16, ηp2=.04), or Time x line x Treatment interaction (F_(1,41)_=.58, p=.45, ηp^2^=.01).

**Figure 7.**
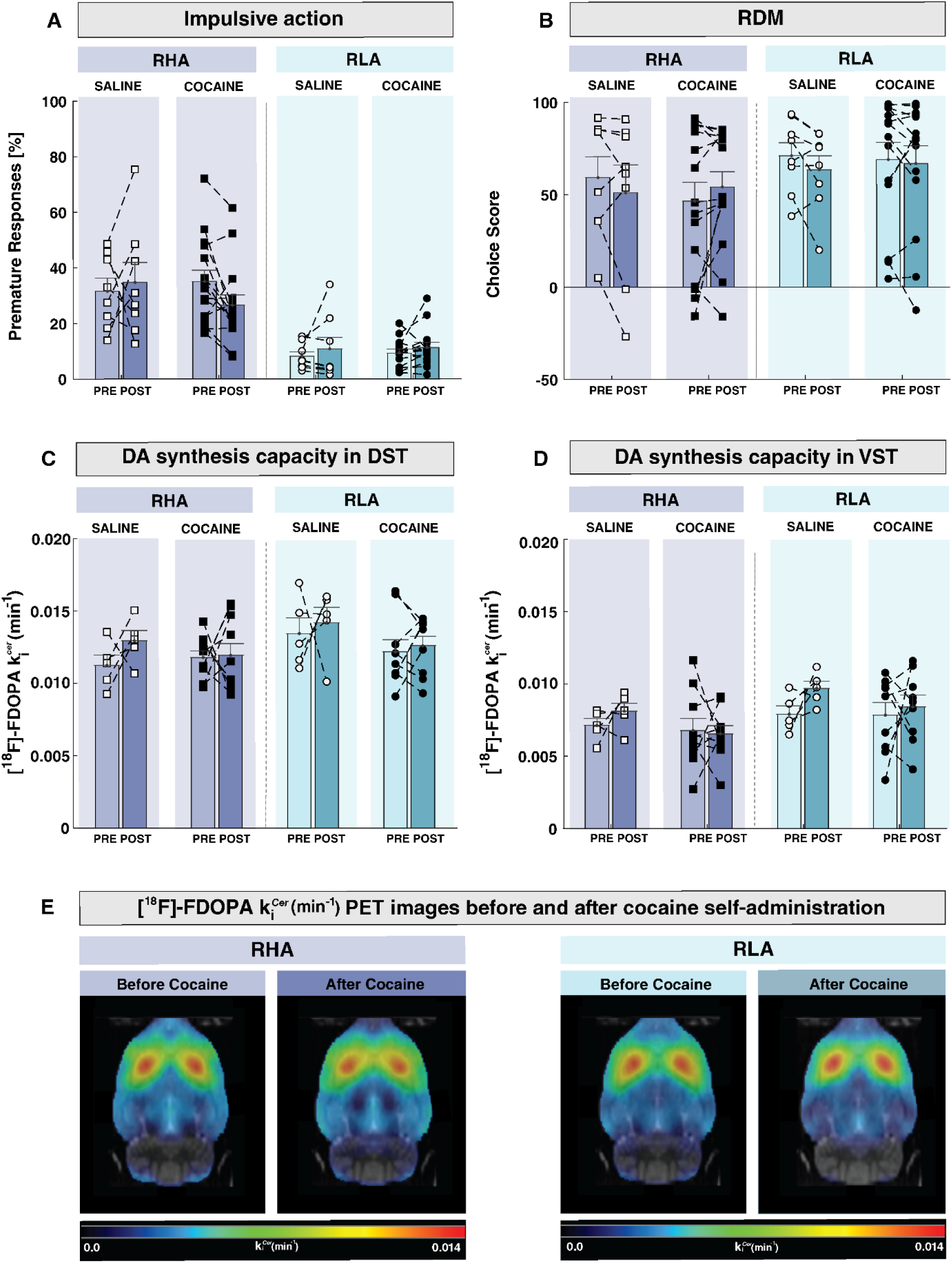
Effect of cocaine SA on impulsive action, RDM, and [^18^F]-FDOPA k_i_ ^cer^ in RHAs and RLAs. Cocaine SA did not alter: **A)** premature responding, or **B)** choice score in RHA or RLA rats. [^18^F]-FDOPA k_i_^cer^, as an index of DA synthesis capacity, was not altered following cocaine SA in either **C)** DST or **D)** VST, suggesting that repeated cocaine intake does not alter DA synthesis. **E)** Averaged parametric maps of[^18^F]-FDOPA k_i_ ^cer^ in RHAs and RLAs before and after cocaine SA. Data are presented as mean ± SEM.

### Effect of repeated Cocaine SA on Striatal DA Synthesis Capacity

When compared with baseline measures, [^18^F]-FDOPA *k_i_ ^cer^* in DST (Fig. 7C) did not significantly change after saline or cocaine SA in any rat line (Treatment: F_(1,25)_=2.7 p=.11, ηp^2^=.09; Treatment x Line: F_(1,25)_=1.4 p=.24, ηp^2^=.05; Time x Line x Treatment interaction: F_(1,25)_=.20 p=.65, ηp^2^=.008). Similarly, an analysis of [^18^F]-FDOPA *k_i_ ^cer^* in VST (Fig. 7D) showed no significant effect of Treatment (F_(1,25)_=3.76 p=.06, ηp^2^=.13), Treatment x Line (F_(1,25)_=.01 p=.91, ηp^2^=0), Time (F_(1,25)_=.06 p=.80, ηp^2^=.002) or Time x Line x Treatment interaction (F_(1,25)_=.0002 p=.99, ηp^2^=0). These results indicate that repeated cocaine taking does not alter baseline DA synthesis capacity in the striatum of RHA or RLA rats (Fig. 7E).

## Discussion

To our knowledge, this is the first longitudinal study in rats investigating the relationship between striatal DA synthesis, impulsive behaviors, and vulnerability to cocaine use. We replicated previous findings showing that RHA rats display higher impulsive action, less optimal decision-making, and higher cocaine-taking than RLA rats (Arrondeau et al., 2023; Bellés et al., 2023; Dimiziani et al., 2019). Importantly, we extended those findings by showing that impulsive action and RDM are not linked to variations in DA synthesis. Additionally, our study shows that chronic cocaine use does not affect impulsive action or RDM, nor decreased DA synthesis, even though similar cocaine exposure has proven sufficient to blunt striatal DA release (Urueña-Mendez et al., 2023). Altogether, our study indicates that alterations in DA synthesis are not the mechanism behind heightened-evoked DA release linked to impulsivity nor the mechanism underlying dopaminergic tolerance to psychostimulants.

Considering that previous studies associated impulsive behaviors with heightened DA release (Bellés et al., 2021) and elevated TH mRNA in the midbrain (Tournier et al., 2013), it is somewhat unexpected that our study found no evidence of altered DA synthesis in impulsive rats. However, a recent study (Nago-Iwashita et al., 2023) showed that TH overexpression does not necessarily result in elevated DA levels, which suggests that additional regulatory mechanisms other than elevated TH expression underly heightened DA release in impulsive subjects. Our findings of preserved DA synthesis in impulsive rats are also consistent with recent neuroimaging studies in humans, where DA synthesis was independent of trait impulsivity (van den Bosch et al., 2023) and RDM (Skandali et al., 2022), despite the relationship between impulsivity and elevated DA release (Buckholtz et al., 2010; Oswald et al., 2015). Our findings are also consistent with microdialysis studies in RHA rats (Giorgi et al., 2003; Lecca et al., 2004) or rats classified as high-impulsive (Zeeb et al., 2016), where basal-extrasynaptic or tonic DA levels were comparable to those of non-impulsive rats. These observations suggest that the heightened evoked DA release linked to impulsive action and RDM does not stem from an overall elevation in DA synthesis or heightened tonic dopaminergic activity, but may more likely arise from an enhanced reactivity of DA neurons to stimulation. Although no studies have directly measured DA cell firing at baseline and in response to stimuli (i.e., rewards or associated cues) in impulsive individuals, the hypothesis that impulsivity might result from altered phasic rather than tonic DA firing has received some indirect support. For instance, optogenetic stimulation of DA neurons in the ventral tegmental area increases premature responding (Flores-Dourojeanni et al., 2021; Suri et al., 2023) and leads to compulsive reward seeking (Seiler et al., 2022). This suggests that impulsive animals might exhibit a heightened reactivity of DA cells to stimulation, which may render them more vulnerable to the rewarding effects of drugs. However, future studies evaluating DA reactivity to rewards and associated cues in animals exhibiting different levels of impulsive behaviors will help to understand whether heightened phasic DA firing underlies impulsivity and the risk of cocaine abuse.

The uncoupling between normal DA synthesis and hyperreactive DA release suggested here is unclear but might result from different mechanisms, including alterations in midbrain D_2_autoreceptor (D_2_AutoR)-inhibitory control of DA neurotransmission. D_2_AutoR negatively controls DA release through mechanisms beyond DA synthesis. For instance, D_2_autoR decreases dopaminergic cell excitability by activating potassium rectifying channels, inhibits DA release by blocking calcium influx, or facilitates DA reuptake by increasing the expression of the DA transporter (Ford, 2014). Hence, a reduction in D2AutoR, as reported in high-impulsive humans (Buckholtz et al., 2010) and rats (Bellés et al., 2021; Besson et al., 2013), could lead to heightened-evoked DA release through any of these mechanisms. Another potential mechanism that might account for an uncoupling between normal DA synthesis and heightened DA release is a deficit in top-down cortical control of DA cell firing. Lesioning or pharmacological inactivation of the medial prefrontal cortex (mPFC) has been shown to elevate impulsive action (Agnoli et al., 2013; Chudasama et al., 2003; Feja and Koch, 2014), RDM (van Holstein and Floresco, 2020; Zeeb et al., 2015), and phasic DA cell firing in the midbrain (Jo and Mizumori, 2016; Tan et al., 2014). Interestingly, in impulsive RHA rats, there is evidence of reduced glutamatergic transmission in the mPFC (Klein et al., 2014) and reduced mPFC volume (Río-Álamos et al., 2019) compared to low-impulsive RLA rats, which could potentially explain their increased impulsivity and hyperreactive DA function. Furthermore, reduced cortical activity has been linked to higher RDM (Lv et al., 2021) and drug abuse in humans (Volkow et al., 1993), and may thus also contribute to a higher vulnerability for cocaine abuse. However, additional experimental evidence is needed to substantiate this hypothesis.

Our results demonstrate for the first time, using a within-subject design, that chronic cocaine exposure does not alter DA synthesis, regardless of baseline levels of impulsivity or the facet of impulsivity assessed. This finding suggests that a reduction in DA synthesis is probably not the mechanism responsible for the attenuation of evoked DA release that is observed in cocaine abusers (Martinez et al., 2007; Volkow et al., 2014) and impulsive animals with a history of cocaine use (Urueña-Mendez et al., 2023). Despite substantial evidence of attenuated DA release in drug abusers (Nutt et al., 2015), studies measuring DA synthesis in this population have been limited, and their results are inconclusive. Consistent with our findings, a study in polydrug users observed no alterations in DA synthesis relative to healthy controls (Tai et al., 2011). Conversely, another study found marked reductions in DA synthesis in cocaine abusers (Wu et al., 1999). Similar contrasts were reported for other drugs, such as cannabis (Bloomfield et al., 2014a) and smoked nicotine (Bloomfield et al., 2014b; Rademacher et al., 2016). In comparison, only a study in rats (Baumann et al., 1993) had previously measured striatal LDOPA accumulation after passive cocaine injections, finding no alterations in exposed animals vs controls. However, it was not clear whether these results were generalizable to self-administered cocaine, which is more representative of drug intake in humans, nor whether the effects varied in impulsive animals. By using within-subjects measures of DA synthesis and a cocaine SA protocol able to blunt striatal DA release in impulsive rats (Urueña-Mendez et al., 2023), our study indicates that reduced DA synthesis does not underlie cocaine-induced DA tolerance, irrespective of baseline impulsivity. Thus, alternative mechanisms might be considered to understand the decreased DA release observed in drug abusers and impulsive rats after chronic cocaine use. For instance, reduced sensitivity of the dopamine transporter (Ferris et al., 2011; Mateo et al., 2005), increased sensitivity or expression of D_2_AutoR (Clare et al., 2021; King et al., 2004; Stefański et al., 2007), or reduced vesicular release (Little et al., 2003; Narendran et al., 2015) have also been observed after chronic cocaine use and could independently lead to reduced capacity of DA release.

Besides exhibiting blunted DA release, cocaine abusers exhibit heightened impulsive behaviors (Kjome et al., 2010; Voon et al., 2014), raising the question of whether impulsivity represents a preexisting risk factor or the consequence of cocaine use. Our data suggest that impulsive action, but not RDM, predicts cocaine-taking, further supporting the view that impulsive action is a stronger predictor than RDM of the rewarding effects of cocaine (Arrondeau et al., 2023) and thus represents an endophenotype of vulnerability to the drug. However, despite continuous research, some debate still exists on the effects of cocaine on specific impulsivity facets. For instance, as shown in our data and consistent with several studies (Abbott et al., 2022; Dalley et al., 2005; Ferland and Winstanley, 2017; Urueña-Mendez et al., 2023), chronic cocaine exposure did not significantly alter impulsive action (but see also: (Caprioli et al., 2013; Dalley et al., 2007; Winstanley et al., 2009). In contrast, the evidence regarding RDM has been less consistent. While previous studies showed impairments in RDM either in risky rats (Ferland and Winstanley, 2017) or in non-risky rats (Cocker et al., 2019), our results show that chronic cocaine use does not alter RDM, irrespective of baseline levels. Although the reason for this discrepancy is unclear, they might reflect differences in the dose of cocaine use. For instance, we used a dose of cocaine that was less than half the dose used in the previous studies. While this dose is sufficient to induce a high response rate in Roman rats (Dimiziani et al., 2019) and trigger DA neuroadaptations under chronic use (Urueña-Mendez et al., 2023), we cannot exclude the possibility that RDM is affected only at higher doses of cocaine.

## Study Limitations

Although RHA rats showed less optimal decision-making than RLA rats, such RDM levels might not be representative of pathological states. Thus, although we observed no alterations in DA synthesis in relation to RDM, DA dynamics might differ in animals with more risky profiles, as suggested by human studies, where heightened RDM in pathological gamblers was linked to increased DA synthesis (van Holst et al., 2018). Another potential limitation of our study is the use of only one temporal window to evaluate the effects of cocaine on DA synthesis. Nevertheless, while cocaine withdrawal might affect DA function (Wu et al., 1999), potential changes in DA synthesis at later withdrawal time are improbable to underlie cocaine-induced tolerance to psychostimulants, as this effect has been observed from early withdrawal (Ferris et al., 2011; Siciliano et al., 2015; Urueña-Mendez et al., 2023). Finally, our study included only males, while the mechanisms controlling DA release may vary in females (Zachry et al., 2021), who are also reported as more vulnerable to cocaine intake than males (Knouse and Briand, 2021). Thus, further studies should consider evaluating sex differences in DA synthesis and its role in impulsivity and cocaine abuse.

**In conclusion, o**ur findings revealed that, unlike DA release (Bellés et al., 2021), striatal DA synthesis does not predict impulsivity or the propensity to cocaine consumption, nor is it altered after chronic cocaine exposure. This suggests that changes in DA synthesis are not the mechanism behind excessive DA release linked to impulsivity, nor the mechanism underlying DA tolerance to psychostimulants. This uncoupling between DA release and synthesis suggests the existence of additional regulatory mechanisms, such as altered DA firing to stimulation, that could contribute to heightened impulsivity and vulnerability to cocaine abuse.

